# The lncRNA *Sweetheart* regulates compensatory cardiac hypertrophy after myocardial injury

**DOI:** 10.1101/2022.11.14.516395

**Authors:** Sandra Rogala, Tamer Ali, Maria-Theodora Melissari, Sandra Währisch, Peggy Schuster, Alexandre Sarre, Thomas Boettger, Eva-Maria Rogg, Jaskiran Kaur, Jaya Krishnan, Stefanie Dimmeler, Samir Ounzain, Thierry Pedrazzini, Bernhard G Herrmann, Phillip Grote

## Abstract

After myocardial infarction in the adult heart the remaining, non-infarcted tissue adapts to compensate the loss of functional tissue. This adaptation requires changes in gene expression networks, which are mostly controlled by transcription regulating proteins. Long non-coding transcripts (lncRNAs) are now recognized for taking part in fine-tuning such gene programs. We identified and characterized the cardiomyocyte specific lncRNA *Sweetheart RNA* (*Swhtr*), an approximately 10 kb long transcript divergently expressed from the cardiac core transcription factor coding gene *Nkx2-5*. We show that *Swhtr* is dispensable for normal heart development and function, but becomes essential for the tissue adaptation process after myocardial infarction. Re-expressing *Swhtr* from an exogenous locus rescues the *Swhtr null* phenotype. Genes depending on *Swhtr* after cardiac stress are significantly occupied, and therefore most likely regulated by NKX2-5. Our results indicate a synergistic role for *Swhtr* and the developmentally essential transcription factor NKX2-5 in tissue adaptation after myocardial injury.

## INTRODUCTION

Precise regulation of gene expression networks is required to form a healthy heart and to maintain proper heart function after birth and throughout adulthood. Such networks not only contain protein coding genes, but also many long non-coding genes (IncRNAs), which are as abundant as coding genes^1,2^. The function and mechanism of these IncRNAs vary greatly, but they are often associated with transcriptional regulation^3^. Two major types of mechanisms are typically discussed when referring to IncRNA gene function. For some loci the resulting RNA is just a circumstantial byproduct with the act of transcription being the major bearer of its function. One such example in the cardiac system is the lncRNA locus *Handsdown RNA* (*Hdnr*), downstream of the *Hand2* transcription factor coding gene. During embryonic development, modification of the *Hdnr* transcriptional activity alters the *Hand2* expression levels, but not modifying the RNA levels of *Hdnr* ^4^. One example of IncRNA loci that exhibit an RNA-based mechanism are the cardiac specific *Myosin Heavy Chain Associated RNA Transcripts* (*MyHEART*; *Mhrt*), a cluster of IncRNAs that undergo anti-sense transcription from the *myosin heavy chain 7* (*Myh7*) locus. Transaortic constriction (TAC) induced pathological stress results in *Mhrt* downregulation by *Brg1* upregulation and subsequent BRG1 mediated chromatin remodeling *in vivo*. Thus, hypertrophy related gene programs are initialized. As the binding of *Mhrt* to BRG1 antagonizes its DNA binding capability, preserving *Mhrt* expression levels after TAC prevents cardiac hypertrophy and heart-failure^5^.

One of the core regulators of heart development is the cardiac specific homeobox protein NKX2-5, which is present in early cardiomyocytes already at embryonic day (E) 7.5 during murine embryonic development. Systemic deletion of the *Nkx2-5* gene in mice causes defects in heart looping and formation of ventricular structures. In addition, other important cardiac regulatory genes are dysregulated and as a combined result the embryos exhibit early embryonic lethality^6^. *Nkx2-5* is not silenced after birth and is abundantly expressed in adult heart tissue^7^. However, not much is known about its function in terminally differentiated cardiac tissue and its involvement in cardiac maintenance and disease. In human patients suffering from congenital heart disease, *Nkx2-5* mutations are commonly found^8^, however, the precise involvement of *Nkx2-5* in the disease context in adult patients remains unknown.

Heart disease represents the main cause of death in the developed world; acute myocardial infarctions (AMI) being the most common form. Reduced blood flow leads to decreased oxygen supply of the heart tissue and thus irreversible damage, such as apoptosis and necrosis of cardiomyocytes and formation of scar tissue which results in a loss of flexibility. Due to the limited regenerative capacity of the terminally differentiated heart tissue, the remaining viable tissue adapts through other mechanisms, such as hypertrophic remodelling that involves thickening of the ventricular walls by an increase of the cardiomyocyte cell size^9^. While cardiac hypertrophy in response to pathological stimuli is often associated with heart failure, we show that it is necessary for survivability after cardiac injury in mice. We characterize a cardiac specific IncRNA we termed *Sweetheart RNA* (*Swhtr*) that is required for regulation of hypertrophic gene programs, most likely acting in concert with NKX2-5.

## RESULTS

### *Swhtr* is a nuclear lncRNA specifically expressed in the heart

In a previously generated dataset that identified the transcriptional landscape of different tissues of early mid-gestation mouse embryos^10^ we identified an RNA that is expressed exclusively in heart tissue. This RNA, which we termed *Sweetheart RNA* (*Swhtr*), is divergently expressed from the essential, transcription factor coding gene *Nkx2-5*^6^. Its annotation partially overlaps with the previously described IncRNA *IRENE-div*^11^. We determined its major transcript by 5’ and 3’ RACE PCR and found that the major variant from *Swhtr* locus is 9,809 nucleotides in length and bears no introns (Fig. 1A). The transcriptional start site (TSS) maps to a previously described GATA4 bound first heart field specific enhancer located approximately 8kb upstream of *Nkx2-5*^12^. To characterize whether *Swhtr* is specific for the first heart field we conducted whole mount *in situ* hybridization (WISH) in E8.25 mouse embryos and found whereas *Nkx2-5* is expressed in the whole heart tube at that stage, *Swhtr* is expressed in the early inflow tract of the developing heart (Fig. 1B). Lineage tracing experiments of *Swhtr* expressing cells confirmed that while *Swhtr* expressing cells contribute to both ventricles, the left loop that originates from the first heart field exhibits a much more even staining (Fig. 1C). The right loop that originates mostly from the second heart field exhibits a more salt-and-pepper like staining (Fig. 1C). In heart and lung of later stage embryos (E12.5; E14.5) the staining within the left and right ventricle is even with no traces of *Swhtr* expressing cells or their descendants found in neither lung nor epicardial tissue (Fig. 1D-E). Compared to the *cis* located *Nkx2-5* gene, *Swhtr* is much lower expressed (Fig. 1A,F). Expression analysis from the whole heart at different stages shows that expression levels of *Nkx2-5* and *Swhtr* are changing comparably (Fig. 1F).

**Figure 1.**
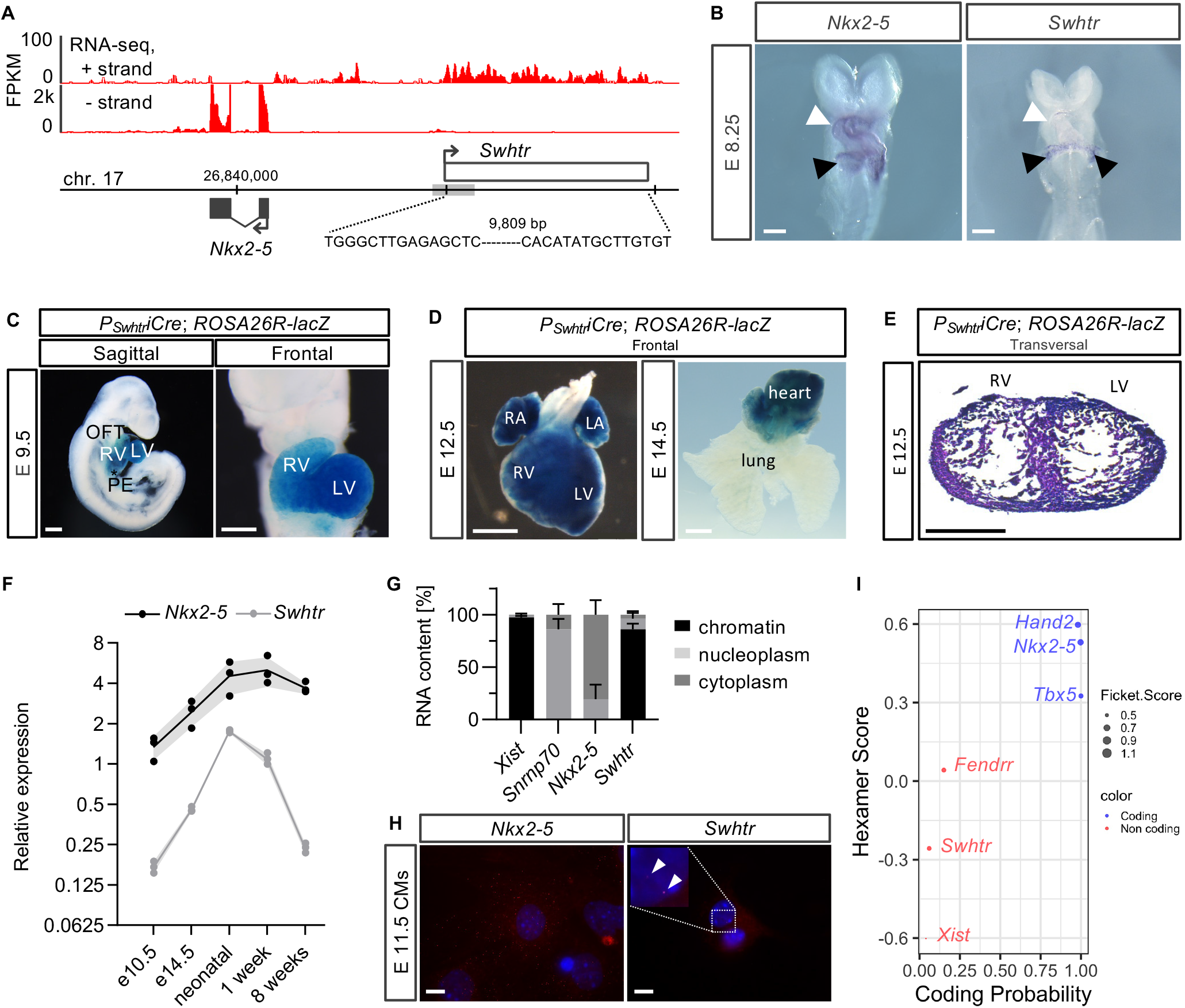
Expression and localization of the *Sweetheart* lncRNA **(A)** Strand-specific RNA-seq from E9.5 heart tubes showing the *Nkx2-5* region. (grey box= first heart field enhancer). Plus-strand track is 20x amplified over the minus-strand track. The number above the genome bar denote the *mm10* co-ordinates and the vertical tip bars represent 10,000 bp steps. **(B)** Whole mount in situ hybridization of *Nkx2-5* and *Swhtr* in E8.25 embryos. White arrow show the early heart tube with two different heart fields and black arrowhead points to the inflow tract region. The white line represents 500 μm. **(C)** Lineage tracing of *Swhtr* expressing cells in E9.5 embryos. OFT= outflow tract, RV= right ventricle, LV= left ventricle, PE= Pre-pericardium. The white line represents 500 μm. **(D)** Lineage tracing of *Swhtr* expressing cells in late gestation embryos and heart/lung explant. The white line represents 1 mm. **(E)** Lineage tracing of *Swhtr* expressing cells in a transversal section of an E12.5 heart. RV= right ventricle, LV= left ventricle. The black line represents 500 μm. **(F)** Quantitative Real-Time PCR timeline of *Swhtr* and *Nkx2-5* expression levels in the hearts of E10.5 embryos to 8 week adult mice. Embryo hearts were pooled from independent litters and the postnatal stages represent data from individual hearts (n=3). **(G)** Subcellular fractionation of E11.5 cardiomyocytes (CMs) of marker transcripts and *Swhtr* (n=2). **(H)** SmFISH of *Nkx2-5* and *Swhtr* in 24h cultured E11.5 cardiomyocytes. The white line represents 10 μm. **(I)** Analysis of coding potential of *Swhtr* by CPAT compared to known coding and non-coding RNAs.

To investigate where the *Swhtr* transcript localizes intracellularly, we conducted subcellular fractionation of embryonic cardiomyocytes. Compared to marker transcripts known to be localized to the chromatin, nucleoplasm and cytoplasm fraction, the *Swhtr* IncRNA localizes predominantly to the chromatin fraction within the nucleus (Fig. 1G). We validated these findings by single molecule fluorescence *in situ* hybridization (smFlSH) experiments that revealed two distinct fluorescent *Swhtr* signals within the nucleus, suggesting that *Swhtr* might reside at its site of transcription (Fig. 1H, Fig. S1). It has become clear, that some RNAs that are classified as lncRNAs can encode micropeptides which might be functional. However, these are usually cytoplasmic localized^13^ pointing against a functional open reading frame (ORF) contained in *Swhtr*. CPAT analysis^14^ further demonstrates that Swhtr has very low coding potential (Fig. 1I). In conjunction with the localization data this points towards a purely non-coding transcript. Together this data shows that *Swhtr* is a chromatin bound IncRNA, specifically expressed in the heart of developing mice and at postnatal stages.

### Genetic inactivation of *Swhtr* does not affect heart development and homeostasis

To investigate the physiological function of *Swhtr* we genetically engineered a knock-out mouse in which we inserted a strong transcriptional stop signal (3xpA) to abolish *Swhtr* expression. To avoid any conflicts with existing regulatory elements we inserted this stop signal downstream of a phylogenetically conserved region of the GATA4 bound enhancer. This genetic insertion causes premature termination of the transcriptional process and, hence, a severely shortened *Swhtr* transcript (Fig. 2A). The full length *Swhtr* RNA is not detectable anymore when we profiled E9.5 hearts by either RNA-seq or qPCR (Fig. 2A,B), demonstrating that the *Swhtr* transcriptional start side (TSS) locates upstream of the inserted transcriptional stop signal and no other alternative transcript is initiated from any downstream elements residing in the *Swhtr* transcription unit. While the *Swhtr* RNA is lost, the expression level of its *cis* located gene *Nkx2-5* in the heart is unchanged at that stage and under these conditions (Fig. 2B).

**Figure 2.**
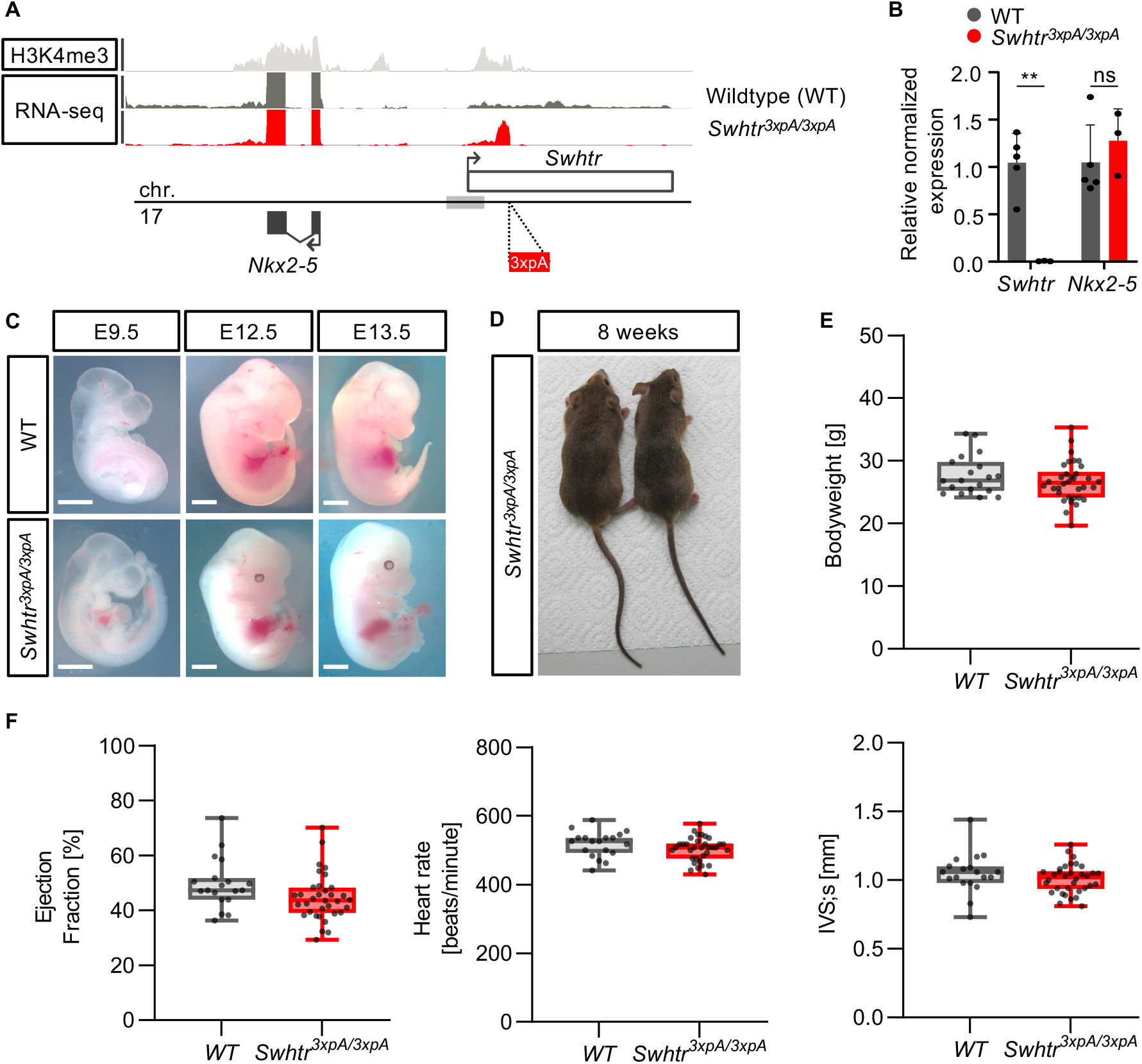
No overt phenotype in *Swhtr*^*3xpA/3xpA*^ mutant embryos or adult mice **(A)** ChIP-seq (H3K4me3) and RNA-seq (E9.5 heart tubes) from WT and *Swhtr*^*3xpA/3xpA*^ mutant mice. **(B)** qPCR validation of loss of *Swhtr* in E9.5 embryonic hearts (WT n=5; *Swhtr*^*3xpA/3xpA*^ n=3). Statistical significance was tested by Two-way ANOVA. ns = not significant, ** < 0.01 **(C)** Embryos of indicated age from WT and *Swhtr* mutants. The white line represents 1 mm. **(D)** Eight week old *Swhtr*^*3xpA/3xpA*^ founder mice. **(E)** Bodyweight of *Swhtr*^*3xpA/3xpA*^ eight times backcrossed (C57Bl6J) mice (WT n=19; *Swhtr*^*3xpA/3xpA*^ n=34). **(F)** Selected cardiac parameters determined by echocardiography in eight week old mice of the indicated genotype (WT n=19; *Swhtr*^*3xpA/3xpA*^ n=34). Statistical significance was tested by Two-way ANOVA. No statistical significant differences were detected.

Phenotypically, homozygous *Swhtr null* embryos do not display differences compared to wild type embryos during development and grow up to adult animals with no overt defects (Fig. 2C,D). To identify subtle phenotypic changes in adult hearts as a result of *Swhtr* lacking throughout development, we investigated the heart function in *Swhtr null* animals by echocardiography after backcrossing the *Swhtr null* mutants to the C57BL/6J genetic background. We compared their body weight and cardiac parameters, such as ejection fraction, heart rate and left ventricular diameter of adult *Swhtr null* and wild type mice. Neither the body weight nor any of the heart parameters differed significantly from that of wild type mice (Fig. 2E-F and Fig. S2). We conclude that under standard breeding conditions *Swhtr* is dispensable for normal heart function.

### *Swhtr* is required for a compensatory response of the heart after myocardial infarction

Many IncRNA knock out animal models do not display an overt phenotype after genetic depletion under standard conditions^15,16,17^. One possibility is that these analyzed lncRNA genes are actually not functional^18^. Another possibility is that a functional requirement is only detected under stress conditions. To challenge the heart we selected the left anterior descending artery (LAD) ligation model, which induces a myocardial infarction in the lateral left ventricle^19^. An acute myocardial infarction (AMI) was induced in male mice of 8 weeks of age and heart parameters were monitored by echocardiography pre-infarction and at day 7 and day 14 after the infarction (Fig. 3A). After 14 days animals were sacrificed, and the presence of infarct tissue was verified by Sirius red staining (Fig. 3G). Notably, compared to wild type mice, *Swhtr null* animals displayed an increased mortality after AMI (Fig. 3B), while the size of the myocardial infarction (MI) was similar between the groups (Fig. S3). Most parameters of cardiac function did not change significantly, but strikingly, the interventricular septum (IVS) of *Swhtr null* mice did not display compensatory thickening after AMI compared to wild type (Fig. 3C-G and Fig. S3). This establishes that *Swhtr* is involved in the adaptive response of the cardiac tissue after a myocardial infarction.

**Figure 3.**
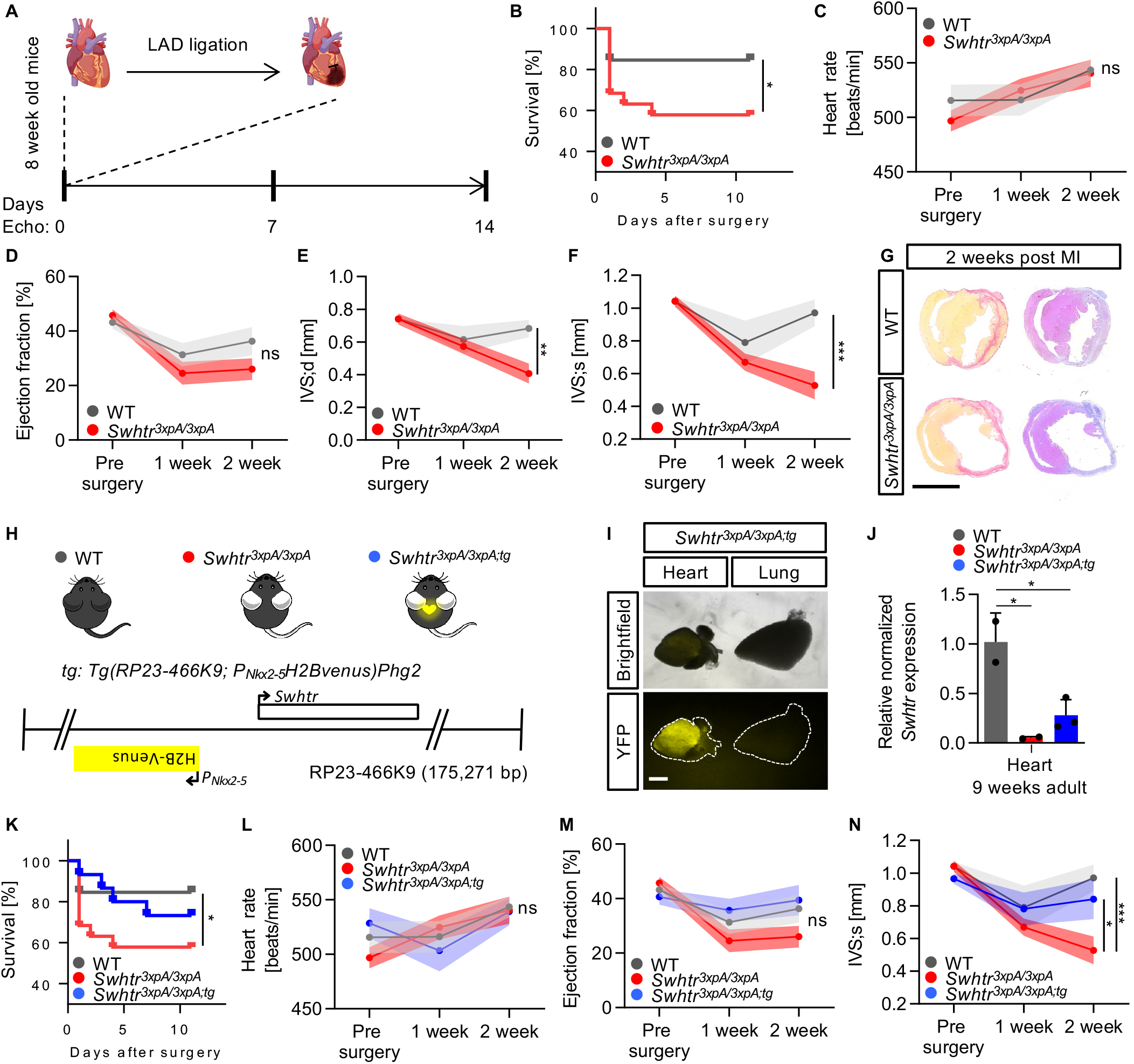
Induced myocardial infarction by left ascending artery ligation (LAD) **(A)** Schematic of of the analysis setup for echocardiography and LAD ligation in mice of age 8 weeks **(B)** Reduced survival of *Swhtr null* mice compared to WT mice after LAD ligation (WT n=13; *Swhtr*^*3xpA/3xpA*^ n=19). Statistical significance was tested by Kaplan-Meier Simple Survival Analysis. * < 0.05 (**C**-**F**) Selected heart specific parameters in mice (n=9 animals per genotype) before and after (1 and 2 weeks) LAD ligation. Statistical significance was tested by Two-way ANOVA. ns = not significant, * < 0.05, ** < 0.01, *** < 0.001 **(G)** Verification of infarct presence in mice 2 weeks after LAD ligation by Sirius red (fibrotic tissue) staining. The black line represents 5 mm. **(H)** Schematic of the rescue transgene and the resulting mouse line. The rescue transgene (*tg*) is comprised of the BAC (RP23-466K9) that includes the *Nkx2-5* (H2Bvenus inserted in *Nkx2-5* ATG) and *Swhtr* loci, randomly inserted into the genome of wild type C57BL6J mice. **(I)** Verification of *tg* presence and activity by heart specific presence of H2BVENUS. **(J)** Expression verification of *Swhtr* in from the *tg* in *Swhtr null* mutants, after crossing. Statistical significance was tested by One-Way ANOVA. * < 0.05 **(K)** Reduced survival of *Swhtr null* mice compared to WT and *Swhtr* rescue mice after LAD ligation (WT n=13; *Swhtr*^*3xpA/3xpA*^ n=19; *Swhtr*^*3xpA/3xpA;tg*^ n=15). Statistical significance was tested by Kaplan-Meier Simple Survival Analysis. * < 0.05 (**L**-**N**) Selected heart specific parameters in mice (n=9 animals per genotype) before and after (1 and 2 weeks) LAD ligation. Statistical significance was tested by Two-way ANOVA. ns = not significant, * < 0.05, *** < 0.001

To determine whether the *Swhtr* dependent adaptation after AMI is a result of the loss of the RNA transcript or the loss of the transcriptional activity at this locus, we generated a *Swhtr* rescue mouse line that re-expresses the *Swhtr* IncRNA from an exogenous locus (random single-copy BAC insertion) (*Tg(RP23-466K9; P*_*Nkx2-5*_*H2Bvenus)Phg2*) (Fig. 3H). This rescue construct contains an *H2BVenus* fusion expression cassette instead of the *Nkx2-5* coding sequence to detect activity of the transgene and to avoid having an additional third copy of the *Nkx2-5* locus present in our genetic setup. Consequently, mice from this transgenic line show yellow fluorescence exclusively in the heart (Fig. 3I) and re-express *Swhtr* (Fig. 3J). We crossed this wild type mouse line to our *Swhtr null* mice to generate the *Swhtr* rescue line (*Swhtr*^*3xpA/3xpA; tg*^).

The *Swhtr* rescue mice do not display the same phenotype as the *Swhtr null* but resemble the wild type control mice in regard to mortality (Fig. 3K) and the size of the IVS (Fig. 3N) after AMI, while neither of the remaining analyzed parameters show significant differences (Fig. 3L-M and Fig. S3). Hence, the RNA of *Swhtr* locus is important for its function.

### *Swhtr* is required for hypertrophic re-modelling after myocardial infarction

The thickening of the interventricular septum after the induced myocardial injury in wild type and *Swhtr* rescue mice might be a result of a hypertrophic response of the remaining muscle tissue. To address whether this might be due to an inherent function of *Swhtr* in cardiomyocytes we first determined the expression of *Swhtr* in different heart cell populations^20^. We found upon fractionation of the four major cell types in the adult heart (8 weeks) that *Swhtr* exhibits the same pattern as the cardiomyocyte marker gene *Tnni1*, showing the cardiomyocyte specificity of *Swhtr* in adult hearts (Fig. 4A). Then, we investigated the presence of larger cells, reminiscent for hypertrophy, in sections of hearts from the LAD-Iigation experiment. Wheat Germ Agglutinin (WGA) staining revealed that the IVS region of wild type and *Swhtr* rescue indeed exhibit on average increased cell size (Fig. 4B,C). In contrast, the cell size in the *Swhtr null* IVS even decreased significantly (Fig. 4C). This establishes that the *Swhtr* RNA is required for the adaptive hypertrophic response of the cardiomyocyte tissue after a myocardial infarction.

**Figure 4.**
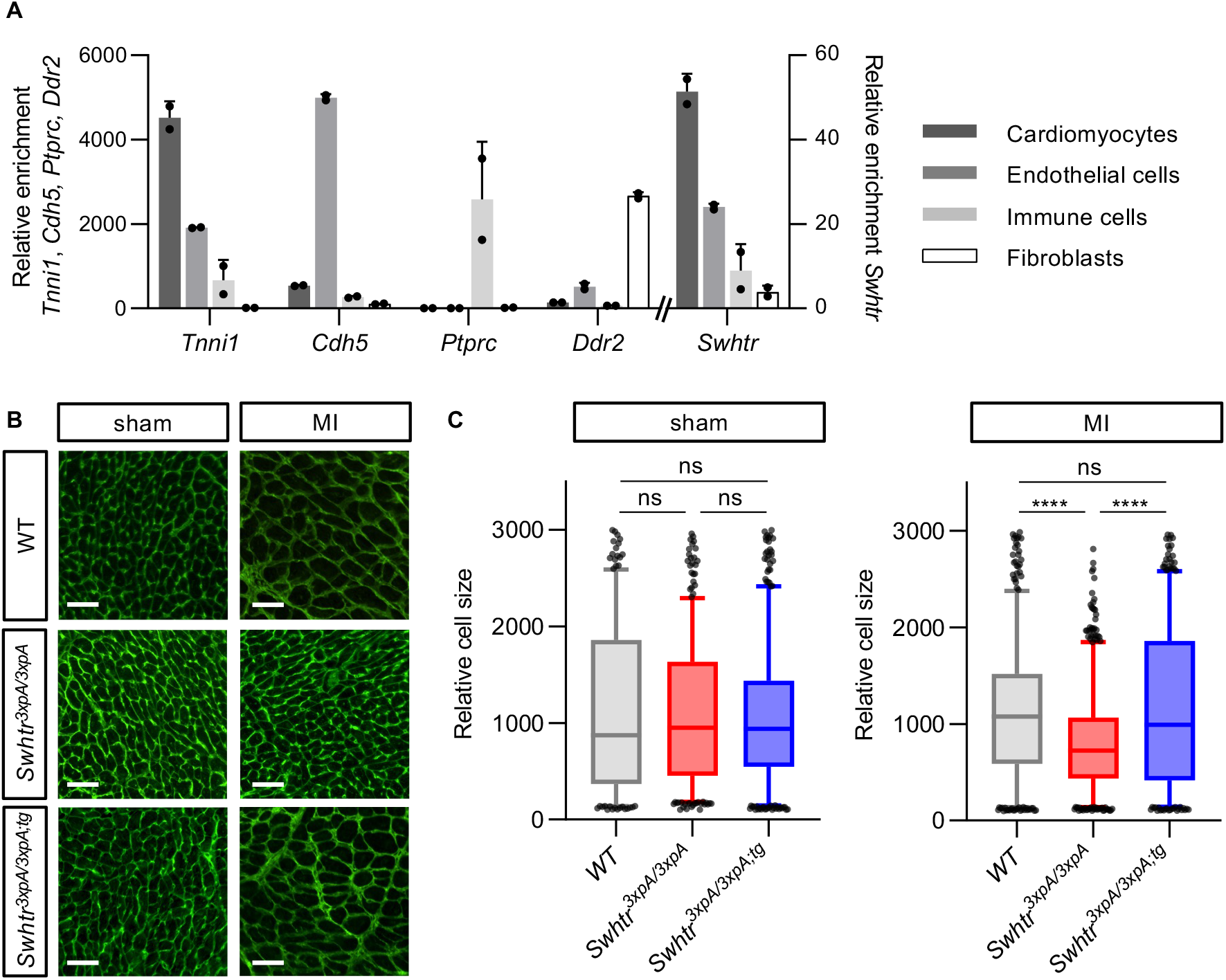
Hypertrophy in *Swhtr* mutant cardiomyocytes. **(A)** Relative enrichment of *Swhtr* in the main cell-types represented in the heart of 8 week old mice as compared to marker genes (n=2). Note that the *Swhtr* expression pattern resembles that of *Tnni1*. **(B)** Wheat Germ Agglutinin stained IHC of representative sections of IVS tissue in the indicated genotypes 2 weeks post sham or MI. The white line represents 50 μm. **(C)** Automated quantifications of relative cell sizes in the interventricular septum of 3 representative animals of the indicated genotype 2 weeks post sham or MI. Statistical significance was tested by One-way ANOVA. ns = not significant, **** < 0.0001

### *Swhtr* regulates NKX2-5 mediated cardiac stress response

Scar formation is one of the consequences of myocardial infarction (Fig. 3E). The resulting heterogeneity of the infarcted heart tissue can lead to biased RNA profiling. To mitigate this compositional bias and obtain consistent RNA profiling we performed primary tissue culture of defined heart tissue slices. Replicate slices were either cultivated continuously for 2 weeks under normoxic condition (untreated) or 1 week under hypoxic condition, followed by 1 week under normoxic condition (treated); mimicking our *in vivo* AMI stress model (Fig. 5A). When we compared expression profiles of wild type heart slices treated versus untreated, we found only 9 genes to be dysregulated (Fig. 5B). In contrast, 464 genes were dysregulated in *Swhtr null* heart slices when treated compared to untreated (Fig. 5B). The near absence of dysregulated genes in wild type compared to the high rate of dysregulated genes in mutant heart slices indicates that *Swhtr* is required for recovery after cardiac stress (Fig. 5B). GO-term analysis shows that these *Swhtr* dependent genes are mainly involved in biological processes such as leukocyte migration and chemotaxis indicative of inflammatory response, intracellular calcium homeostasis, muscle function, heart morphogenesis and extracellular matrix organization, all of which are integral components of cardiac stress response (Fig. 5C). Accordingly, KEGG pathway analysis reveals pathways involved in inflammatory response, cardiomyopathy, glucose metabolism and response to oxygen levels depend on *Swhtr* (Fig. S4). Albeit the loss of *Swhtr* does not lead to changes in *Nkx2*-*5* expression level neither in adult hearts (Fig. 2B) nor in our treated heart slice culture (Fig. 5B), the most likely localization of *Swhtr* to its locus of transcription (Fig. 1I) implicates some involvement of *Nkx2-5* to this process. To determine if a significant number of *Swhtr* dependent genes are occupied by NKX2-5 and therefore might be direct targets we analyzed available CHIP-seq data of NKX2-5 occupation in 6 weeks old adult heart tissue^21^. We found that a significant number of dysregulated genes is occupied by NKX2-5 (Fig. 5D), indicating that the lack of *Swhtr* impairs NKX2-5 mediated response to cardiac stress. This suggests that *Swhtr* might act together with NKX2-5 to regulate this cardiac stress response. One possibility is that at some timepoint during the response to cardiac stress stimuli *Swhtr* might modulate *Nkx2-5* expression levels to regulate its transcriptional effect on its target genes in adult cardiomyocytes.

**Figure 5.**
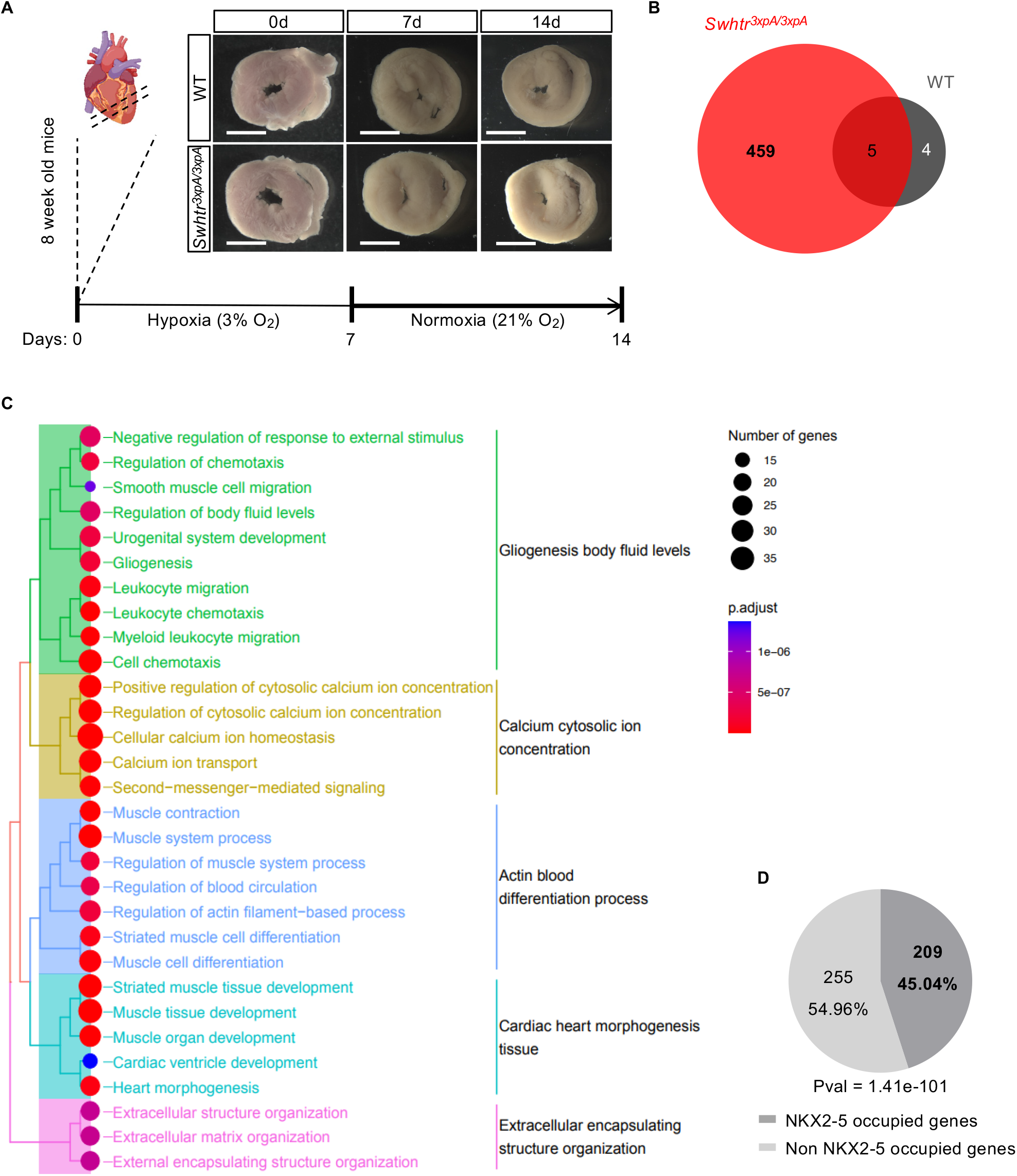
*Swhtr* dependent genes under cardiac stress **(A)** Schematic overview of the experimental procedure with representative pictures of heart slices. The white line represents 5 mm. **(B)** Number of deregulated genes after 7 days of hypoxia treatment followed by 7 days of normoxic conditions of *Swhtr*^*3xpA/3xpA*^ heart slices compared to WT heart slices (n=4). **(C)** GO-term enrichment analysis of deregulated *Swhtr* dependent genes. **(D)** NKX2-5 occupation on *Swhtr* dependent genes. P-value from hypergeometric test (ChIPpeakAnno).

## DISCUSSION

Here we characterize a novel IncRNA that we termed *Swhtr*, which partially overlaps with a previously published lncRNA from that locus: *IRENE-div*^11^. In contrast to this previously described eRNAs, our study shows that the absence of the long transcript from the *Swhtr* locus does not lead to persistent *Nkx2-5* mRNA dysregulation, neither during embryonic development nor cardiac homeostasis. We show by fractionation and smFISH that the *Swhtr* lncRNA is chromatin localized, arguing together with the high non-coding potential that no micropeptide is embedded in the nearly 10kb long *Swhtr* transcript. Our data clearly show an RNA based function for *Swhtr*, as rescue animals do not exhibit the same defect in cardiac hypertrophy as the *Swhtr null* animals. The promoter of *Swhtr* was described previously as a *Nkx2-5* cardiac enhancer element active in multipotent cardiac progenitor cells^22^. The *Swhtr* promoter becomes active after myocardial stress, which was described previously^12^. It was tested if these reappearing cells can contribute to regeneration of an infarcted heart, but our data rather indicate that this genetic element is important to activate *Swhtr* and that this activity is required for the tissue remodeling of the remaining intact cardiac tissue after infarction.

The *cis* located gene to *Swhtr* is the core cardiogenic transcription factor coding gene *Nkx2-5. Nkx2-5* is known to be required for embryonic development of the cardiac system^6^. However, although the locus is still active and abundantly expressed in adults, not much is known about its function after birth and in cardiac stress. Interfering with NKX2-5 function by over-expression of a dominant negative *Nkx2-5* mutant has been shown to lead to apoptosis of cultured cardiomyocytes. In contrast, over-expression of a wild type form of *Nkx2-5* has a protective effect against doxorubicin induced stress^23^. This demonstrates a critical role of *Nkx2-5* in maintenance of adult cardiomyocytes as well as in the response to stress. Here, we show significant NKX2-5 occupation on the *Swhtr* dependent genes. This indicates that *Swhtr* acts in concert with NKX2-5 under stress conditions to regulate hypertrophy associated gene programs. Mutations in *NKX2-5* have been associated previously with dilated cardiomyopathy^24^, but we show for the first time that *Nkx2-5* and its wider locus is an important responder to cardiac stress in adult hearts *in vivo*, acting in concert with its divergently expressed lncRNA *Swhtr*.

In response to increased demands on the remaining tissue after injury, the viable tissue adapts by cardiac hypertrophy to maintain blood supply to the body^9^. This is demonstrated by the increase of average cell size in the IVS of wild type animals after AMI, together with the heart parameters being unaffected. Our *Swhtr null* mice have defects in this hypertrophic response after cardiac injury. While the differences of ejection fraction do not reach statistical significance, the observed decrease meets the requirements of a trend in loss of heart functionality. Together with the increase of lethality after myocardial injury this points towards a cardioprotective role of the *Swhtr* lncRNA by making the cardiac tissue permissive for a hypertrophic response after myocardial injury. This is further supported by our *ex vivo* analysis of heart slices. One of the main pathways known to be involved in hypertrophic process is the phosphoinositide 3-kinase (PI3-K) pathway that is, among others, responsible for metabolic substrate utilization and function of cardiomyocytes^25^. The RNA profiling of cardiac slices subjected to hypoxic stress and following recovery time, revealed dysregulation of 464 genes in slices derived from *Swhtr null* animals as compared to only 9 dysregulated genes in wild type slices. Notably, KEGG pathway enrichment analysis revealed that AGE-RAGE signaling is among the most affected pathways. Even though it is not frequently discussed in relation to cardiac hypertrophy, it is a pathway known to be involved in stress response of cardiomyocytes to stimuli such as oxidative stress^26,27^). Additionally, AGE-RAGE signaling is known to be an inducer of the PI3-K pathway, coherent with its role in hypertrophic remodeling.

Maladaptive hypertrophy, in contrast, is defined as occurring after pathological stimuli and associated with heart failure and disease^28^. Among others, inflammatory response, calcium homeostasis and signaling, and extracellular matrix deposition are biological processes involved in maladaptive cardiac hypertrophy^29^. These biological processes are significantly impaired upon loss of *Swhtr*, indicative for maladaptive behavior. It has been discussed that an initial adaptive hypertrophic response can transition to maladaptive hypertrophy upon peresistant pathological stress^29^. Within the scope of the experiment, no decrease of heart function could be detected in animals capable of hypertrophic remodeling, despite the deregulation of maladaptive hypertrophic processes in dependence of *Swhtr*. Additionally, increased survival of wild type animals compared to *Swhtr* mutant animals suggests that hypertrophic remodeling has a positive effect on viability after acute myocardial infarction and is a required compensatory response.

While the detailed mechanism remains to be determined we here show a clear cardioprotective role of the murine lncRNA *Swhtr* and a compensatory gene regulatory network depending on *Swhtr*. While no lncRNA was described yet for the human *NKX2-5* locus, the promoter element of *Swhtr* is highly conserved across placental species. Moreover, in public datasets from human heart tissue some RNA seems to be present around this conserved promoter element. Further investigation can determine whether this *SWHTR* locus might have the same cardioprotective role in humans and if this function could be employed for a therapeutic application.

## Acknowledgements

We thank Dijana Micic and Sonja Banko for excellent animal husbandry and Karol Macura for the generation of the transgenic mice. This research was supported by the DFG (German Research Foundation) Excellence Cluster Cardio-Pulmonary System (Exc147-2). T.A and S.R. are supported by the 403584255 – TRR 267 of the DFG.

## Data availability

The data are deposited to GEO and can be downloaded under the accession number GSE200380. The cDNA of *Sweetheart RNA* (*Swhtr*) is deposited with GenBank under ON351017.

## EXPERIMENTAL PROCEDURES

### Culturing of mouse ES cells

The genetic background of the ES cells generated in this work is identical (129S6/C57BL6 (G4))^30^ or C57BL/6J (gift from Lars Wittler). The mESCs were either cultured in feeder free 2i media or on feeder cells (mitomycin inactivated SWISS embryonic fibroblasts) containing LIF1 (1000 U/ml). 2i media: 1:1 Neurobasal (Gibco #21103049):F12/DMEM (Gibco #12634-010), 2 mM L-glutamine (Gibco), 1x Penicillin/Streptomycin (100x penicillin (5000 U/ml,) /streptomycin (5000ug/ml), Sigma #P4458-100ML, 2 mM glutamine (100x GlutaMAX™ Supplement, Gibco #35050-038), 1x non-essential amino acids (100x MEM NEAA, Gibco #11140-035), 1x Sodium pyruvate (100x, Gibco, #11360-039), 0.5x B-27 supplement, serum-free (Gibco # 17504-044), 0.5x N-2 supplement (Gibco # 17502-048), Glycogen synthase kinase 3 Inhibitor (GSK-Inhibitor, Sigma, # SML1046-25MG), MAP-Kinase Inhibitor (MEK-Inhibitor Sigma, #PZ0162), 1000 U/ml Murine_Leukemia_Inhibitory_Factor ESGRO (10^7^ LIF, Chemicon #ESG1107), ES-Serum media: Knockout Dulbecco’ s Modified Eagle‘s Medium (DEMEM Gibco#10829-018), ES cell tested fetal calf serum (FCS), 2 mM glutamine, 1x Penicillin/Streptomycin, 1x non-essential amino acids, 110 nM ß-Mercaptoethanol, 1x nucleoside (100x Chemicon #ES-008D), 1000 U/ml LIF1.

The cells were split with TrypLE Express (Thermo Fisher Scientific #12605-010) and the reaction was stopped with the same amount of Phosphate-Buffered Saline (PBS Gibco #100100239) followed by centrifugation at 1000 rpm for 5min. The cells were frozen in the appropriate media containing 10% Dimethyl sulfoxide (DMSO, Sigma Aldrich #D5879). To minimize any effect of the 2i^31^ on the developmental potential mESC were only kept in 2i for the antibiotic selection for transgene integration after selection kept on feeders.

### Genetic manipulation of ES cells and generation of embryos and mice

ES cells were modified according to standard procedures. Briefly, 10×10^6^ ES cells were electroporated with 25μg of linearized targeting construct and cultivated with selection media containing 250 μg/ml G418 (Life Technologies #10131035) or 125 μg/ml Hygromycin B (Life Technologies #10687010) for the first and second targeting, respectively. Resistant clones were isolated, and successful gene targeting was confirmed. Embryos and live animals were generated by tetraploid complementation^32^. Homozygous *Swhtr*^*3xpA[N]/3xpA[H]*^ ES cells generated 21 mice from four foster mothers, confirming their integrity and usability in subsequent developmental assays. The selection cassettes consisting of *PGK::Neo-SV40pA* (abbreviated “N”) or *PGK::Hygro-SV40pA* (abbreviated “H”) were flanked by *FRT* sites. Selection cassette was removed by crossing animals with a FLP delete strain^33^.

### Generation of *Swhtr* Rescue BAC

The BAC RP23-466K9 was ordered from BACPAC Resource Center (BPRC) and its integrity verified by HindIII digest. The RP23-466K9 BAC contains the 175,271bp of mouse genomic sequence surrounding the *Nkx2-5* locus. This BAC includes the upstream (*Bnip1*) and the downstream (*Kifc5b*) located genes. The BAC was modified using the Red/ET recombinase system (Genebridges). The H2B-Venus expression cassette, followed by a bglobin polyadenylation site and a downstream Neomycin selection cassette, was inserted into the ATG of *Nkx2-5*. This will eliminate any *Nkx2-5* expression from the BAC and simultaneously allows monitoring of rescue construct by means of detection of yellow fluorescence in the hearts of embryos and adults.

Around 3 Mio mESC cells of the C57Bl6J background were collected and resuspend in 680 μl PBS and were mixed with 120 μl linearized (PI-SceI) BAC (42 ng/μl). The BAC was electroporated into the C57Cl6J cells under the following conditions: 240V; 500uF; 4mm; ∞ with a Gene Pulser Xcell™ Electroporation Systems from BioRad. Afterwards the cells were resuspended in 2i Media and plated on gelatin coated cell culture dishes. The next day the selection of the cells started with 300 μg/ml G418 (InvivoGen, #ant-gn-1). The selection was done till the colonies were big enough for picking after 7-8 days. Afterwards the procedure was the same as described above.

### Generation of mouse embryos and strains from mESCs

All animal line generation procedures were conducted as approved the Landesamt für Gesundheit und Soziales Berlin (LAGeSo), Berlin under the license numbers G0349/13. Embryos were generated by tetraploid morula aggregation of embryonic stem cells as described in^30^. SWISS mice were used for either wild-type donor (to generate tetraploid morula) or transgenic recipient host (as foster mothers for transgenic mutant embryos). All transgenic embryos and mESC lines were on a hybrid F1G4 (C57Bl6/129S6) background or the C57Bl6J background (rescue BAC).

To generate the mouse strains the transgenic cells were aggregated with diploid morula SWISS embryos. The genotype of the cells was either hybrid F1 for the *Swhtr* mutant mice or wild type C57BL6J for the rescue mice. Adult *Swhtr* mutant mice were backcrossed 6 times to C57BL6J before all subsequently conducted experiments.

### Whole mount *in situ* hybridization

Whole-mount *in situ* hybridization was carried out using standard procedures described on the MAMEP website (http://mamep.molgen.mpg.de/index.php). Probes were generated by PCR from E11.5 heart ventricle cDNA using primer containing promotor binding site for T7 and SP6 polymerase. After verification of the probe templates, antisense *in situ* probes were generated as described on the MAMEP website using T7 polymerase (Promega #P2077). The *in situ* probes are generated against *Nkx2-5 or Sweetheart*.

### RNA isolation

To isolate RNA either from heart tissue or cultivated cardiomyocytes the cells were lysed in 900 μl Qiazol (Qiagen, #79306). To remove the DNA 100 μl gDNA Eliminator solution was added and 180 μl Chloroform (AppliChem, #A3633) to separate the phases. The extraction mixture was centrifuge at full speed, 4°C for 15min. The aqueous phase was mixed with the same amount of 70 % Ethanol and transferred to a micro or mini columns depending of the amount of tissue and cells. The following steps were done according to the manufactural protocol.

### Subcellular RNA fractionation

Cellular fractionation was carried out as previously described^34^. Briefly, cell pellets were resuspended in 200 μl cold cytoplasmic lysis buffer (0.15% NP-40, 10mM Tris pH 7.5, 150mM NaCl) using wide orifice tips and incubated on ice for 5 min. The lysate was layered onto 500 μl cold sucrose buffer (10mM Tris pH7.5, 150mM NaCl, 24% sucrose w/v), and centrifuged in microfuge tubes at 13,000 rpm for 10 min at 4 C. The supernatant from this spin (700 μL) represented the cytoplasmic fraction. 10% (70 μL) of the supernatant volume was added to an equal volume of 2X sample buffer for immunoblot analysis. The remaining supernatant was quickly added to 15 ml tubes containing 3.5X volumes of QIAGEN RLT Buffer, supplemented with 0.143 M ß-mercaptoethanol. RNA purification from these and subsequent cellular fractions was performed according to manufacturer instruction.

The nuclear pellet was gently resuspended into 200 μl cold glycerol buffer (20 mM Tris pH 7.9, 75 mM NaCl, 0.5 mM EDTA, 50% glycerol, 0.85 mM DTT) using wide orifice tips. An additional 200 μl of cold nuclei lysis buffer (20 mM HEPES pH 7.6, 7.5 mM MgCl2, 0.2 mM EDTA, 0.3M NaCl, 1M urea, 1% NP-40, 1mM DTT) was added to the samples, followed by a pulsed vortexing and incubation on ice for 1 min. Samples were then spun in microfuge tubes for 2 min at 14,000 rpm and at 4 C. The supernatant from this spin represented the nucleoplasmic fraction (400 μL), and 10% of supernatant was kept for immunoblot analysis. 3.5X volumes of QIAGEN RLT were added to the remaining nucleoplasmic supernatant.

50 μl of cold PBS was added to the remaining chromatin pellet, and gently pipetted up and down over the pellet, followed by a brief vortex. The chromatin pellet was extremely viscous and sticky, and therefore difficult to fully resuspend. 5 ml of the PBS supernatant was collected for immunoblot analysis as above, and 500 μl TRI-Reagent was added to the pellet. After vigorous vortexing to resuspend the chromatin, chromatin-associated RNA was extracted by adding 100 μl chloroform and incubated at room temperature for 5 min. The chromatin samples were then centrifuged in microfuge tubes for 15 min at 13,000 rpm at 4 C. The resulting upper aqueous layer was then added to 3.5X volumes QIAGEN RLT buffer.

### Full-length cDNA determination

Rapid amplification of cDNA end (RACE) was performed using the SMARTer® RACE 5’/3’ Kit (Takara, #634858). 1ug of freshly isolated RNA from E8.5 embryo hearts was used to generate first strand cDNA according to the manufactural protocol. The primers were designed between 60-70°C Tm. Half of the PCR product was analysed by agarose gel electrophoresis and the rest was used for nested PCRs to validate the 5’ and 3’end. PCR products were extracted from the agarose gel and sent for sequencing. After the determination of the end, the full-length sequences were amplified and sequenced. The sequences were deposited at GeneBank under the IDs: ON351017 (*Sweetheart RNA, Swhtr*).

### Fractionation of the main cell types of the adult heart

Fractionation was conducted as previously described^20^. Briefly, adult mice were sacrificed and hearts collected in HBSS (gibco #14025050). Hearts were enzymatically digested using the Multi Tissue Dissociation Kit 2 (Miltenyi #130-110-203). Cardiomyocyte fraction was obtained by pre-plating. Endothelial cells and immune cells were obtained by magnetic separation. Fibroblasts were enriched by another preplating step.

### Cardiac Injury Models – Ligation of the left anterior descending artery

Mouse was anesthetized by IP injection of a mixture of ketamin/xylazine/acepromazin (65/15/2 mg/kg). Mouse was placed on warming pad for maintenance of body temperature. In the supine position, endotracheal intubation was performed, and the mouse was placed on artificial ventilation with a mini-rodent ventilator (tidal volume = 0.3ml, rate = 120 breaths/min). Ocular gel was applied to hydrate the cornea during the surgical procedure. Proper intubation was confirmed by observation of chest expansion and retraction during ventilated breaths. A left thoracotomy was performed. The pectoralis muscle groups were separated transversely, and the fourth intercostal space was entered using scissors and blunt dissection. The pericardium was gently opened, and a pressure was applied to the right thorax to displace the heart leftward. A 7.0 silk ligature was placed near the insertion of the left auricular appendage and tied around the left descending coronary artery. Occlusion of the artery was verified by the rapid blanching of the left ventricle. The lungs were re-expanded using positive pressure at end expiration and the chest and skin incision were closed respectively with 6-0 and 5-0 silk sutures. The mouse was gradually weaned from the respirator. Once spontaneous respiration resumed, the endotracheal tube was removed, and the animal was replaced in his cage on a warming pad with standard chow and water ad libitum. Analgesic drug (Temgesic, Buprenorphin 0.1 mg/kg) was administered subcutaneously after the surgery.

Animal experiments were approved by the Government Veterinary Office (Lausanne, Switzerland) and performed according to the University of Lausanne Medical School institutional guidelines.

### In vivo transthoracic ultrasound imaging

Transthoracic echocardiography was performed using a 30 MHz probe and the Vevo 2100 Ultrasound machine (VisualSonics, Toronto, ON, Canada). Mice were lightly anesthetized with 1-1.5% isoflurane, maintaining heart rate at 400-500 beats per minute. The mice were placed in decubitus dorsal on a heated 37ºC platform to maintain body temperature. A topical depilatory agent was used to remove the hair and ultrasound gel was used as a coupling medium between the transducer and the skin. The heart was imaged in the 2D mode in the parasternal long-axis view. From this view, an M-mode cursor was positioned perpendicular to the interventricular septum and the posterior wall of the left ventricle at the level of the papillary muscles. Diastolic and systolic interventricular septum (IVS;d and IVS;s), diastolic and systolic left ventricular posterior wall thickness (LVPW;d and LVPW;s), and left ventricular internal end-diastolic and end-systolic chamber (LVID;d and LVID;s) dimensions were measured. The measurements were taken in three separate M-mode images and averaged. Left ventricular fractional shortening (%FS) and ejection fraction (%EF) were also calculated. Fractional shortening was assessed from M-mode based on the percentage changes of left ventricular end-diastolic and end-systolic diameters. %EF is derived from the formula of (LV vol;d – LV vol;s)/LV vol;d*100. Echographies were done in baseline condition and one and two weeks after surgery. Sacrifices were done the day of the 2-week post-MI echography.

### Heart preparation and histology

Adult hearts were dissected two weeks after MI and fixed in 4% paraformaldehyde/PBS over night. Fixed hearts were embedded in paraffin and sections (4-6 μm thickness) were mounted onto Superfrost® Plus microscope slides (Thermo scientific #630-0950). Immunohistochemistry was carried out using standard procedures. The Antibody used for the detection of cell borders was anti wheat germ agglutinin (WGA, Thermo Fisher Scientific, #W11261). The slides were mounted with Vectashield (VWR, #101098-042) and sealed with colorless nail polish. Image documentation was conducted using the NIKON Eclipse Ci, equipped with the Ds-Ri2 color camera. Analysis of cell sizes was conducted using ImageJ software.

### Real-time quantitative PCR analysis

Quantitative PCR (qPCR) analysis was carried out on a StepOnePlus™ Real-Time PCR System (Life Technologies) using Power SYBR® Green PCR Master Mix (Promega #A6002). RNA levels were normalized to housekeeping gene. Quantification was calculated using the ΔΔCt method^35^. *Hmbs* served as housekeeping control gene for qPCR. The primer concentration for a single reaction was 250nM. Error bars indicate the standard error from biological replicates, each consisting of technical duplicates. A list of oligonucleotides can be found in Table S1.

### Embryo / heart preparation and histology

Staged embryos and adult hearts were dissected from uteri into PBS and fixed in fresh 4% paraformaldehyde/PBS 1mm tissue per 1h at 4°C. For histology, embryos and hearts were embedded in paraffin. E7.5 embryos were removed from the uterus of timed mated mothers together with the surrounding decidua and fixed all together. Sections (4-6 μm thickness) were mounted onto Superfrost® Plus microscope slides (Thermo scientific) or on Zeiss MembraneSlide 1.0 PEN NF (#415190-9081-001). The stainings were carried out with Eosin (Carl Roth), Hematoxylin (AppliChem) and Sirius Red according to standard procedures. All image documentation was carried out on Microscope Leica M205C with the MC170 HD camera and captured with ImageJ. Except the E7.5 embryo sections, which were imaged on a NIKON Eclipse Ci, equipped with the Ds-Ri2 color camera.

### SmFISH

FISH Probes were designed, using the biosearchtech.com/stellaris-designer website and ordered from BioCat. The lyophilized Probes were resuspended in 400 μl 1x Tris-EDTA Buffer (10 mM Tris-HCl (ApliChem, #A1086, ApliChem, #A5634), 1 mM EDTA (Life technologies, #15575020), pH 8.0) to get a final concentration of 500nM per μl. The probes were conjugated with a Quasar570 dye and small aliquots were stored at -20°C.

Cardiomyocytes from E11.5 heart ventricles were cultivated (as described) on cover slips (10 mm Marienfeld, #0111500) for 48h. For the fixation process the cells were washed once with PBS and fixed 10min at room temperature with 4%PFA/PBS (AppliChem, #A3813). Again, the cells were washed 3 times with PBS and permeabilized 5 min on ice with permeabilize sol (1xPBS, 1%RNAse inhibitor Ribovanadylcomplex (RVC, NEB,#S1402S), 0,5 % Triton X-100 (Sigma, #T8787)). Afterwards the cells were washed three times with 70 % Ethanol (Roth, #T913.7) and stored in 70 % Ethanol at -20°C. For the hybridization transfer the cover slips to 70% Ethanol at room temperature and incubate 10min. Add wash buffer (10% saline sodium citrate buffer (20xSSC-Buffer, Invitrogen, # 15557-036), 10% Formaldehyde (FA, Sigma, # F8775), in RNAse free water) and incubate again for 10min. Afterwards incubate the cells with 25 μl of hybridization solution (10 % SSC-Buffer, 10 % FA, 10% Dextran sulfate (Roth, # 5956.3), in RNAse free water, 2ul of the dye (1000nM)) in a humidity chamber at 37°C in the dark for 4h. Transfer the cells to pre-warmed wash buffer and incubate in the dark at 37°C 30min without shaking. Afterwards wash the cover slips with 2xSSC Buffer and incubate 5min at room temperature. Mount the cover slips with Vectashield Mounting Media containing DAPI (VWR, # 101098-044) and seal it with colorless nail polish. For the visualization Zeiss Axio Observer-Z1 with a 100x objective was used.

### Heart slice preparation, treatment and harvest

4 Wild-type and 4 *Swhtr*^*3xpA/3xpA*^ mice were sacrificed by cervical dislocation when they reached the age of 8 weeks. The heart was collected in HBSS and washed in BDM buffer (HBSS + 10 mM BDM) to remove blood cells. Apex and base of the heart were manually removed before preparing the slices using a sharp scalpel. The slices were washed in BDM buffer once more before submerging them in culture medium consisting of DMEM (gibco#10569010) supplemented with 10 % Fetal Bovine Serum, 1% Non-Essential Amino Acids (gibco #11140050) and 1% Penicillin-Streptomycin (gibco #15140122). The heart slices were left to recover overnight. During the experiment, the media was refreshed every other day.

For the treatment, the slices were incubated in a humidified hypoxic chamber (3% O_2_, 5% CO_2_) at 37 °C for 7 days and moved to a humidified incubator with normoxic conditions (21% O_2_, 5% CO_2_) at 37 °C for 7 additional days. Slices that were incubated in a humidified incubator at normoxic conditions (21% O_2_, 5% CO_2_) at 37 °C for 14 days served as the control. Following the treatment, the slices were harvested in 1 ml TRI reagent (Sigma-Aldrich #T9424) and homogenized using Precellys® 2 mL Soft Tissue Homogenizing Ceramic Beads. RNA was extracted by Phenol/Chloroform extraction. Briefly, 200 μl Chloroform were added per 1 ml of TRI reagent and mixed. After centrifugation at 4 °C aquous phase was precipitated by adding 0.7 vol of Isopropanol and incubating at –20 °C for 1 hour. The precipitated RNA was pelleted by centrifugation at 4 °C, washed with 70 % Ethanol and resuspended in nuclease-free water.

### Bioinformatic analysis and data deposit

RNA was treated to deplete rRNA using Ribo-Minus technology. Libraries were prepared from purified RNA using ScriptSeq™ v2 and were sequenced on an Illumina novaseq 6000 platform at Novogene. We obtained 25 million paired-end reads of 150 bp length. Read mapping was done with STAR aligner using default settings with the option --outSAMtype BAM SortedByCoordinate^36^ with default settings. For known transcript models we used GRCm38.102 Ensembl annotations downloaded from Ensembl repository^37^. Counting reads over gene model was carried out using GenomicFeatures Bioconductor package^38^. The data are deposited to GEO under the accession number GSE200380.

All genes with read counts < 10 were excluded. For normalization of read counts and identification of differentially expressed genes we used DESeq2 with Padj < 0.05 and log2FC= 0.58 cutoff^39^. GO term were analyzed using clusterProfiler and enrichplot Bioconductor packages^40^. To overlap NKX2-5 binding with *Swhtr* dependent DE genes, we downloaded ChIP-seq data from E12.5 heart (GSM3518650). Raw reads were downloaded and aligned to mm10 using Bowtie2^41^. Samtools^42^ was used to convert aligned reads to sorted bam files. Duplicated reads as well as reads overlapping blacklisted region were removed using bedtools^43^. Peaks, then, were called with MACS3 peak caller^44^. All peaks were sorted, merged and finally intersected with genes coordinates using bedtools and ChIPpeakAnno Bioconductor package^45^. The pvalue of the overlapping peaks were calculated according to hypergeometric test.

