## Supplementary material for "The lncRNA *Sweetheart* regulates compensatory cardiac hypertrophy after myocardial injury": S1 S2 S3 S4

**Rogala, Supplementary Figure 1**

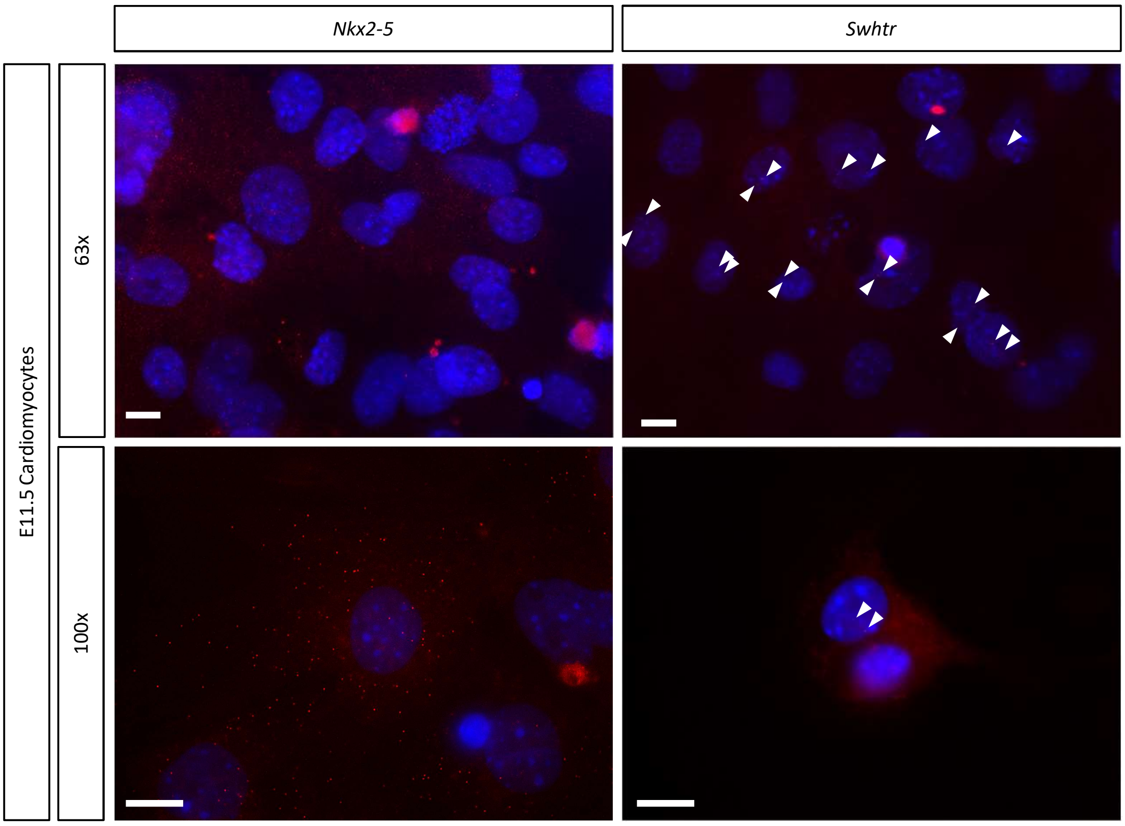


**Supplementary Figure 1**, smFISH of *Nkx2-5* and *Swhtr* on E11.5 primary cardiomyocytes
Two different magnifications are shown. Arrowheads indicate signals and the white line represents 10 µm.

**Rogala, Supplementary Figure 2**

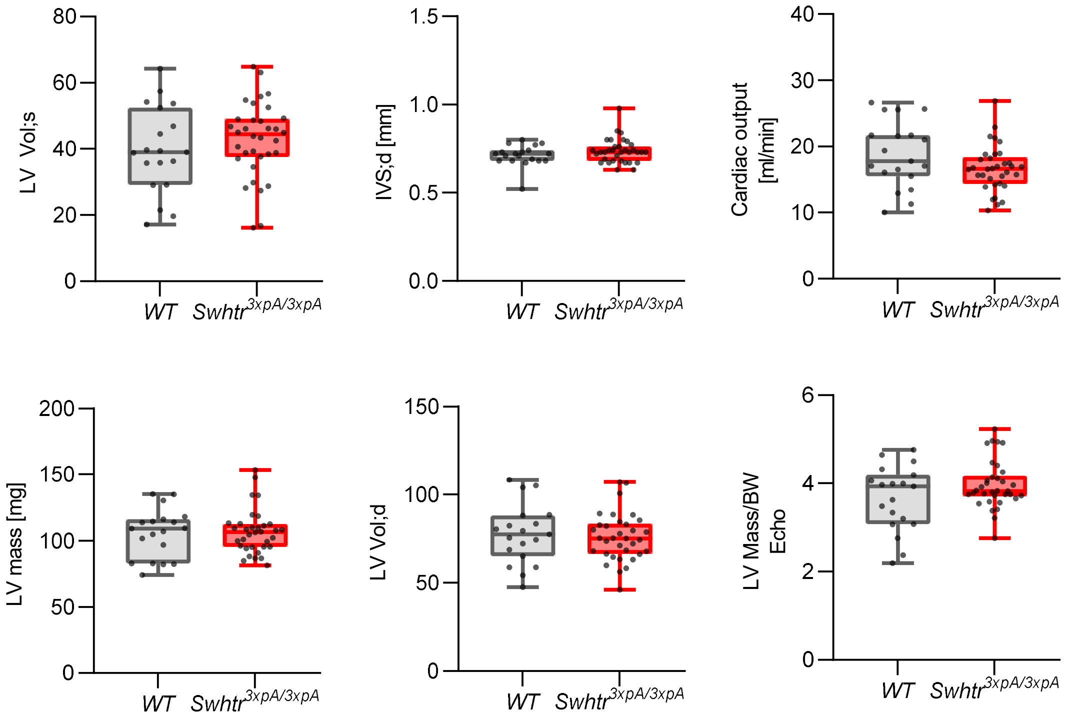


**Supplementary Figure 2**, No difference in heart parameters in *Swhtr*$^{3xpA/3xpA}$ mutant mice Statistical significance was tested by Two-way ANOVA. No statistically significant differences were detected.

**Rogala, Supplementary Figure 3**

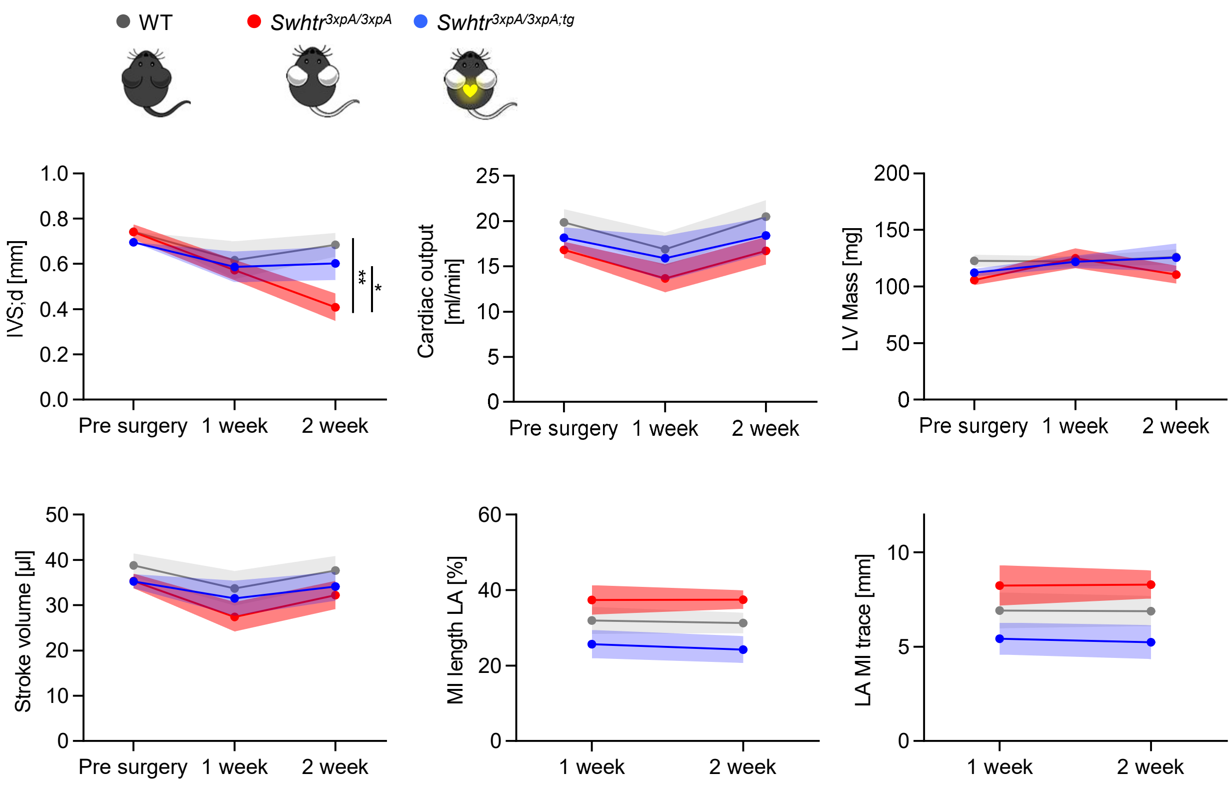


**Supplementary Figure 3**, Additional heart parameters after MI
Statistical significance tested by Two-way ANOVA. No indication implies that no statistically significant difference from the wild type was detected. * < 0.05, ** < 0.01 (n=9 animals per genotype)

**Rogala, Supplementary Figure 4**

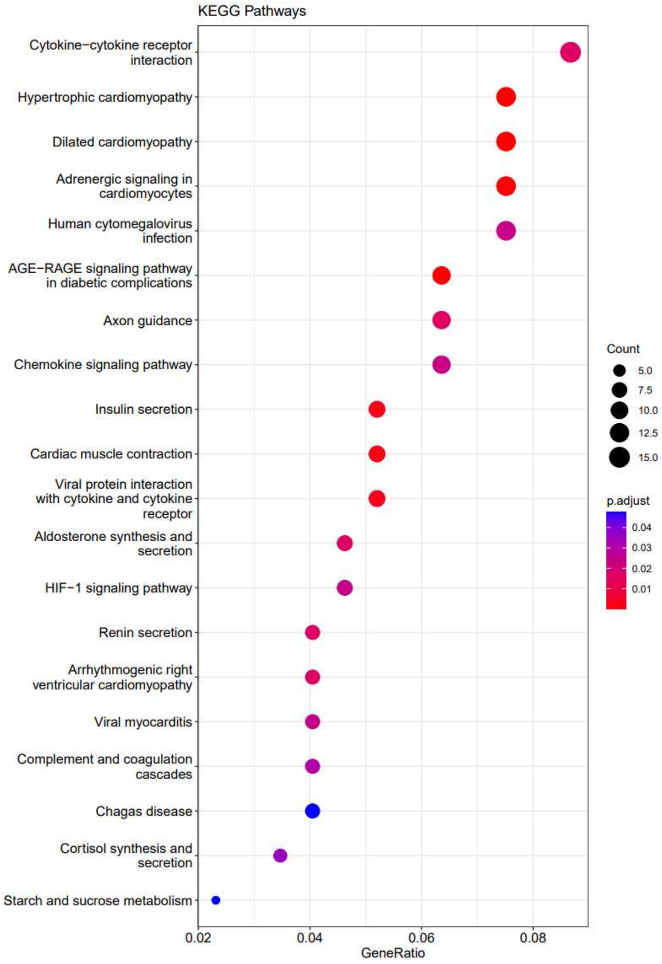


**Supplementary Figure 4**, KEGG-pathway-analysis of *Swhtr* dependent genes
