## Supplementary material for "The lncRNA *Sweetheart* regulates compensatory cardiac hypertrophy after myocardial injury": Table S1

**Table S1 – related to Materials & Methods**

**Oligos and probes used for indicated experiments**

| qPCR analysis | Forward | Reverse |
|---|---|---|
| *Nkx2-5* | AAGTGCTCTCCTGCTTTCCCA | TTTGTCCAGCTCCACTGCCTT |
| *Swhtr* | CCGCAAACCGAACAACTCAG | CTGCCTAAGTGGGGGAAGTG |
| *Eaf1* | agagctgtggtgttggcatt | agagagcaaaacctggagcc |
| *Tnni1* | CCATGGATCTGCGGGCCAACC | GCGGCCTTCCATGCCAGACA |
| *Cdh5* | TTGGACCGAGAGAAACAGGC | CTTGCCCACTCGGATGTCTT |
| *Ptprc* | GGCTTCAAGGAACCCAGGAA | GAGTGCCTTCCTCCATGCTT |
| *Ddr2* | AGCGAGGTACAGGACTCCAT | TCCCCGCTCCTCCTCAATTA |
| *Xist* | agggtgtgtgtgcatatgga | ccgccatcttttcctgtacg |
| *Snrnp70* | tgctcctcctccaacaagag | cagagtcttgaaggcatcgc |
| *Hmbs* | CCTGGGCGGAGTCATGTC | ACTCGAATCACCCTCATCTTTGA |

| RACE analysis | Reverse |
|---|---|
| 5´ *Swhtr* | GATTACGCCAAGCTTTCTCCATCCCGCTTTTCACC |
| 3´ *Swhtr* | GATTACGCCAAGCTTATCCCCTGGAACTGGGGTTA |

| smFISH probes | *Nkx2-5* | *Swhtr* |
|---|---|---|
| 1 | ctggagtagggggggattcag | accacttacaacctgattca |
| 2 | caggtgggtagcagagagtg | atgtgcaccttgaaagcttg |
| 3 | gagaaaggcgtgggtgtgag | aaatgctccttttaagggct |
| 4 | caggttcaggatgtctttga | gttcacactaattggtgtgc |
| 5 | gaaagcaggagagcacttgg | aaaggccctatcgattactt |
| 6 | taggctcccgggtaaaatgt | gaagtggagcagttgaagca |
| 7 | gcgcacagctcttttttatc | caaagagtatctgggcctac |
| 8 | ctgcgagaagagcacgcgtg | gtattgcagccaagaagtga |
| 9 | ggtaccgctgttgcttgaag | ggcaattggaagggagcaag |
| 10 | tggacgtgagcttcagcacg | ctcagcagtttgagtaatcc |
| 11 | tggaaccagatcttgacctg | ctactttgtttccgtacaga |
| 12 | tcgcttgcacttgtagcgac | aacctttgctattgccaag |
| 13 | agaagctccagagtctggtc | cctgaaggaaggactagtt |
| 14 | ttgtagccataggcattgag | ctgaacactctgcctaagtg |
| 15 | gacgccaaagttcacgaagt | tgttagagggaaagacaccc |
| 16 | gactctgcacggtgttcaag | taggacaggaagagagccat |
| 17 | ttctcctaaaggtgggagtc | gaaaaggtctaagtgtccct |
| 18 | tgggtgtgaaatctgaggga | ttaccaaccaggcactatag |

| | | |
|---|---|---|
| 19 | gggatggatcggagaaaggt | gtcacagagatagagcagga |
| 20 | acaccaggctacgtcaataa | gtgttctttcgttcagtttg |
| 21 | gtgtggaatccgtcgaaagt | ctagttttcgggtcaagagt |
| 22 | cgacggcaagacaaccaggg | tcatgaacccacattgcata |
| 23 | aacataaatacgggtgggtg | gtcaagaatgctcaacctgg |
| 24 | gagggttctcatttcttaca | gtttcagactgtaggcattt |
| 25 | gttagcgcactcactttaat | ctttccatttgataggcaga |
| 26 | | gggatacttgcctacatatt |
| 27 | | tatccctttggattttgagt |
| 28 | | ccatccagaatggatagagt |
| 29 | | aaatccatagcattggctgg |
| 30 | | atagctcgattctctaggtg |
| 31 | | caacctcaaggccagaagaa |
| 32 | | agactttccttggttgaagg |
| 33 | | aaatcgcttagactgcgtgg |
| 34 | | gacccaaaacaacagccatg |
| 35 | | ctagttcttaccaccaagta |
| 36 | | gcccaatagttagaacatgc |
| 37 | | ctagaaaccaaagccctact |
| 38 | | ttgtcctgtggacttaacac |
| 39 | | tcctggtgctcaaggtaaaa |
| 40 | | gatgtccaggatggaaatgc |
| 41 | | aatacattggaacaccctgc |
| 42 | | ttccaatctgtgcagaagtc |
| 43 | | aggcccaggtaaaagaactc |
| 44 | | ttgttatctattgtgccacg |
| 45 | | atgcattactatatagccca |
| 46 | | attcacatttatcctgacct |
| 47 | | tatcttattccatgctcact |
| 48 | | tgccaactgaaccagtttta |

| Genotyping | Forward | Reverse | Internal |
|---|---|---|---|
| *Swhtr3xpA/3xpA* | CCCGCATGAAGATTCTGGGA | GCCTGAAATGAGCCTTGGGA | CAGGTTGGAAGTGGAGCAG |
| *Swhtr3xpA/3xpA;tg* | TGACCGAGTACAAGCCCAC | GCGCAGACAGGTCCCCAGAC | |

| WISH probes | Forward | Reverse |
|---|---|---|
| *Nkx2-5* | aATTTAGGTGACACTATAGAA<br>CAAGTGCTCTCCTGCTTTCC | aTAATACGACTCACTATAGG<br>GTGGAATCCGTCGAAAGT |

| | | |
|---|---|---|
| *Swthr* | aATTTAGGTGACACTATAGAA<br>AATCTTTGGGCCAGGACTTT | aTAATACGACTCACTATAGG<br>TCCACCTCTTTTTCCATTGC |
